# Energetic failure and oxidative stress underlie the *Prpf31* splicing factor-related mouse phenotype

**DOI:** 10.1101/2025.02.28.640751

**Authors:** Florian Hamieh, Elora M Vanoni, Quentin Rieu, Julie Enderlin, Nawel Hadjout, Déborah S. Lew, Anaïs Potey, Géraldine Millet-Puel, Thierry Léveillard, Emeline F. Nandrot

**Author notes:** These authors contributed equally to this work (Florian Hamieh, Elora M Vanoni). **<u>Corresponding author:</u>** Emeline F. Nandrot, Institut de la Vision, 17 rue Moreau, Paris, F-75012, France.

## Abstract

Mutations of ubiquitous *PRPF* splicing factors represent the second cause of retina-specific autosomic dominant retinitis pigmentosa. *Prpf31* downregulation decreases phagocytosis of mouse and human retinal pigment epithelial (RPE) cells, thus suggesting similar pathogenesis between species. With time, the mouse RPE ultrastructure shows signs of cellular stress such as cytoplasmic vacuoles. To decipher the primary cellular origin of *Prpf31*-related deleterious processes we first confirmed the gradual accumulation of protein and lipid oxidations. We then showed deregulation in the expression levels of oxidative and endoplasmic reticulum stress markers as well as of mitochondrial respiratory chain constituants, first and foremost in the RPE from 3 months onward. For the first time we analyzed the energetic metabolism of freshly dissected RPE/choroid, retina and peritoneal macrophages, and showed that mitochondrial respiration and global energy production were decreased solely in *Prpf31^+/-^* RPE cells. Therefore, our results indicate that metabolic impairments and associated stress might contribute to pathogenesis first in *Prpf31*^+/-^ RPE cells before affecting the retina.

## Introduction

Retinitis pigmentosa (RP) is the most common form of inherited retinal degeneration affecting around 1:4,000 individuals worldwide. It is an heterogeneous disease at both the clinical and genetic levels characterized by a progressive vision loss due to photoreceptors degeneration associated with a visual field constriction and preceeding complete blindness (Hartong et al., 2006). Mutation of the *pre-mRNA processing factors* (*PRPF*) family of splicing factor genes represents the second most frequent cause of autosomic dominant RP (adRP, 15-20% of cases) (Audo et al., 2010; Hartong et al., 2006; Mordes et al., 2006; Ruzickova and Stanek, 2017; Van Cauwenbergh et al., 2017; Wheway et al., 2020). These ubiquitous proteins are involved in the assembly of the U4/U6.U5 tri-snRNP complex of the spliceosome and crucial for proper transcript splicing (Grainger and Beggs, 2005; Makarova et al., 2002). Interestingly, not all patients with these mutations do develop the RP phenotype. This heterogeneity is explained by an haploinsufficiency linked to wildtype alleles with weaker expression –due to nonsense mediated decay– and to variation in the number of a minisatellite repeat elements, thus resulting in incomplete penetrance and suggesting a “dose-related” phenotype (Maubaret et al., 2011; Rio Frio et al., 2008; Rivolta et al., 2006; Rose and Bhattacharya, 2016; Rose et al., 2016; Tanackovic et al., 2011; Vithana et al., 2003). Some cases have also been associated with dominant negative and gain-of-function effects (Mordes et al., 2006; Wheway et al., 2020).

Strikingly, mutations of these ubiquitously expressed genes give solely rise to a retina-specific phenotype, both in patients and animal models (Chakarova et al., 2002; Farkas et al., 2014; Graziotto et al., 2011; Hamieh and Nandrot, 2019; Hebbar et al., 2021; Linder et al., 2011; McKie et al., 2001; Vithana et al., 2001; Yin et al., 2011). Hence, understanding how ubiquitous proteins lead to such a restricted phenotype has been an endeavor taken by several research groups. One characteristic of these factors, especially for Prpf31, is its higher expression after birth in the retina and even more in retinal pigment epithelial (RPE) cells facing photoreceptors, when compared to other tissues (Cao et al., 2011; Valdés-Sanchez et al., 2019; Yuan et al., 2005). Indeed, these ocular tissues have high requirements in RNA processing and protein synthesis for their daily functions. Some *PRPF31* mutations have been shown to potentially hinder the translocation of the protein to the nucleus and making the protein unstable, rendering splicing insufficient especially in high demand conditions (Deery et al., 2002; Huranová et al., 2009; Towns et al., 2010; Valdés-Sanchez et al., 2019; Wilkie et al., 2006). Studies in drosophila, zebrafish and Human-derived retinal organoids suggested that photoreceptors were primarily affected by degeneration (Azizzadeh Pormehr et al., 2020; Hebbar et al., 2021; Linder et al., 2011; Yin et al., 2011). While photoreceptor-related genes have been associated with PRPF31-containing complexes (Mordes et al., 2007), other studies highlighted the potential link of splicing factors with cilia-related phenotypes (Buskin et al., 2018; Maxwell et al., 2021; Wheway et al., 2015). However, retinal dystrophies can be complex regarding pathogenesis, and photoreceptor degeneration is sometimes a secondary effect linked to mutations in RPE cells (Hartong et al., 2006).

We previously reported the characterization of mouse models harboring mutations in the *Prpf3* (*Prpf3^T494M^*), *Prpf8* (*Prpf8^H2309P^*) and *Prpf31* (*Prpf31^+/-^*) genes, *Prpf31^+/-^* mice being maintained as heterozygotes due to embryonic lethality (Bujakowska et al., 2009; Farkas et al., 2014; Graziotto et al., 2011). Surprisingly, no typical RP phenotype was observed in these strains beside a diminished rod function in old *Prpf3^+/T494M^* mice (Graziotto et al., 2011). However, a more recent mouse model with a tissue-specific homozygous *Prpf31* mutant allele generated using the CRISPR/Cas9 approach confirmed that total absence of Prpf31 in the retina leads to photoreceptor degeneration in mice like in Human patients (Xi et al., 2022). At the cellular level, we and other laboratories detected a similar ≈40% decrease in retinal phagocytosis of RPE cells from *Prpf31*^+/-^ primary cultures, the Human ARPE-19 cell line down-regulated for *PRPF31* as well as *PRPF31*-RP11 patient-derived RPE from induced pluripotent stem (iPSC-RPE) cells (Brydon et al., 2019; Buskin et al., 2018; Farkas et al., 2014; Rodrigues et al., 2022). Despite the difference in pathological timing between patients and mouse models possibly due to differences in the splicing machineries between species or between diurnal and nocturnal species, these results suggest that common mechanisms occur in both species and that mouse models can be used as a paradigm to identify related pathological processes at the molecular level.

Indeed, these mice are affected at different functional levels pointing to potential molecular targets of Prpf deficiency. In fact, as early as 2 months of age we detected defects in the *in vivo* rhythmic regulation of two RPE crucial functions, the phagocytosis of photoreceptor outer segments (POS) and retinal adhesion (Farkas et al., 2014). With age, we found accumulation of morphological changes in RPE cells: loss of basal infoldings, formation of amorphous deposits beneath the RPE and presence of vacuoles in the cytoplasm, all suggestive of stress processes (Graziotto et al., 2011; Valdés-Sanchez et al., 2019).

Aging organisms in normal conditions undergo oxidative stress, mitochondrial dysfunction and accumulation of damaged proteins (Gray et al., 2003; Scarpulla, 2008; Serrano and Klann, 2004; Trojanowski and Mattson, 2003). These different types of stress can be exacerbated in pathological situations and produce cumulated cellular dysfunctions as it was largely described in other retinal diseases as well as in RPE cells (Kaarniranta et al., 2020; Masuda et al., 2017; Markitantova and Simirskii, 2025). In addition, stress of the endoplasmic reticulum (ER) and the associated unfolded protein response (UPR) have been associated with several types of retinal degenerations (Zhang et al., 2014). Thus, we set out to characterize RPE-related stress pathways in *Prpf31^+/-^* mice at different ages. We analyzed oxidative and ER stress, detoxification pathways, mitochondrial respiration and other energy-producing metabolic pathways in 3- to 18-month-old animals in order to dissect the full series of molecular events. We also studied if the combination of oxidative stress and Prpf31 deregulation had an even more deleterious effect on RPE phagocytosis.

## Results

### Prpf31 deficiency increases oxidative stress and impairs detoxification pathways

Based on our previous study, RPE from *Prpf31^+/-^* mice display abnormal apical and basement membrane structures and accumulation of cytoplasmic vacuoles with age (Graziotto et al., 2011). These results suggested that mutant RPE cells could be subjected to stress conditions, we therefore checked for oxidation of DNA, proteins and lipids. Increased levels of nitrotyrosine immunostaining in RPE cells and of 4-hydroxynonenal (4-HNE) labeling in RPE and photoreceptors have been detected at 18 months (Fig. 1 A). The abundance of these markers suggests high levels of reactive oxygen species (ROS) at the protein and lipids levels in *Prpf31^+/-^* mice compared to wildtype (wt) littermate controls.

**Figure 1.**
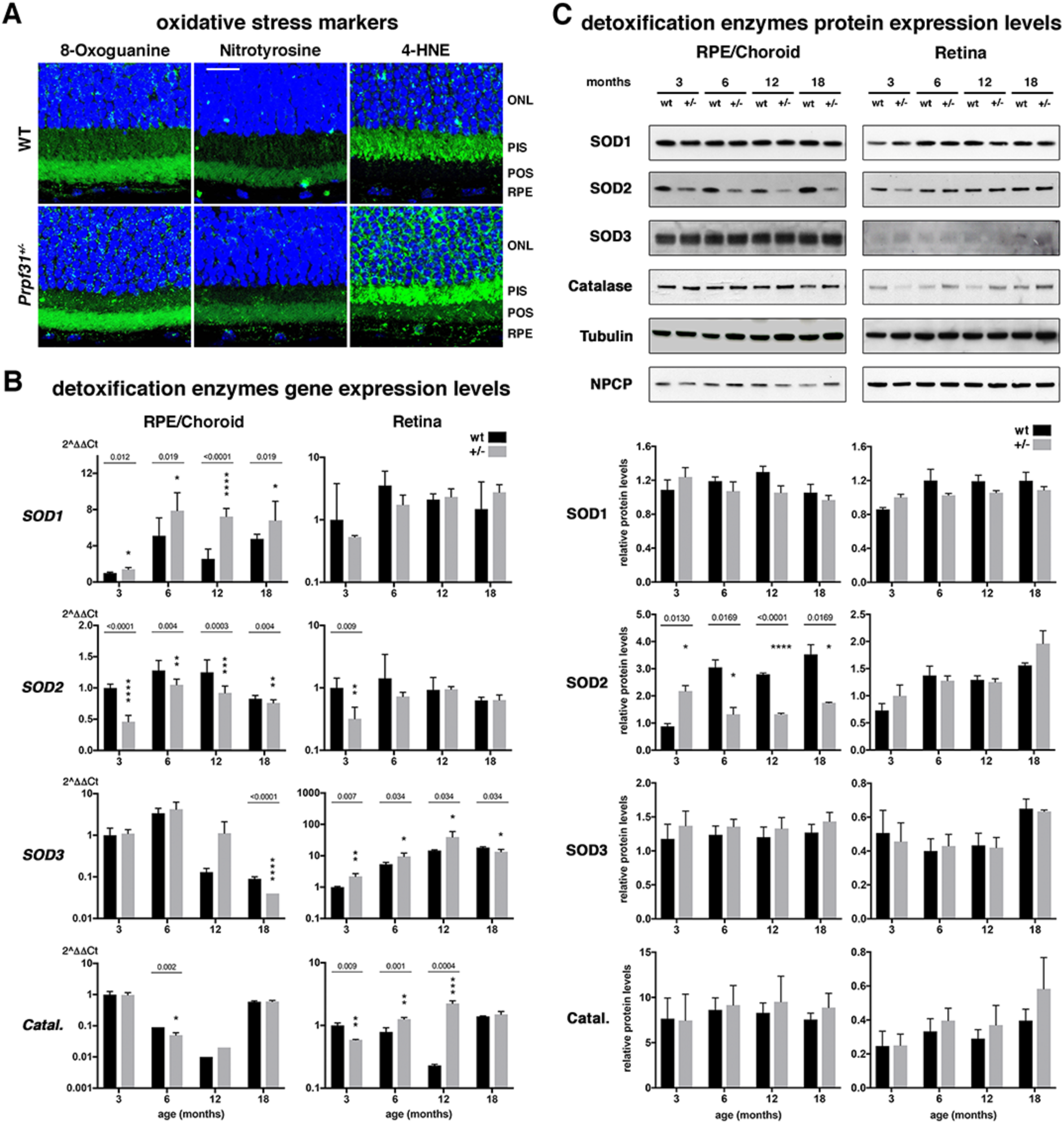
Oxidative stress and deregulated detoxifing enzymes in RPE/choroid and retina tissues from *Prpf31^+/-^* mice. **(A)** Confocal immunohistochemistry images of wildtype (WT) and *Prpf31^+/-^*retinal sections for 8-oxoguanin (left), nitrotyrosine (center) and 4-hydroxynonenal (right) oxidative stress markers (green). Nuclei are labeled with DAPI (blue). Scale bar 20µm. RT-qPCR **(B)** and immunoblot **(C)** expression profiles of SOD1, SOD2, SOD3 and catalase detoxifing enzymes in wildtype (wt) and *Prpf31^+/-^* (+/-) RPE/choroid and retina fractions at different ages (months) as indicated. Tubulin and NPCP were used as internal loading controls for immunoblots. Data expressed as 2^ΔDCt relative to wt control samples at 3 months (B) or as relative protein levels compared to a control sample (C). Immunoblot band sizes (B): SOD1 20kDa, SOD2 25kDa, SOD3 35kDa, catalase 65kDa, tubulin 60kDa, NPCP 70kDa. Mean ± s.d., n = 3-12 (B, except n = 2 for *Catalase* in *Prpf31^+/-^* RPE/choroids at 6 and 12 months and in wt retinas at 3 months) and n = 3-4 (C), multiple *t* test with significance threshold values set as * *P* < 0.05, ** *P* < 0.01, *** *P* < 0.001, **** *P* < 0.0001.

To investigate the efficiency of intracellular detoxification systems activated in presence of oxidative stress, mRNA and protein expression levels of superoxide dismutases (SOD) SOD1, SOD2, SOD3 and catalase were analyzed at different ages. The gene encoding cytoplasmic Cu/Zn *SOD1* was significantly overexpressed in *Prpf31^+/-^*mice’s RPE/choroid but not in retinal fractions at all ages, but this was not translated at the protein level whatever the age or genotype (Fig. 1 B, C). Mitochondrial Mn *SOD2* was downregulated in *Prpf31^+/-^* RPE/choroid fractions at all ages, but no real difference was observed in corresponding retinal samples. SOD2 protein levels were increased in mutant samples at 3 months but increased in older animals consistently with RNA samples. A slight upregulation in the expression of *SOD3* and *Catalase* were observed only in *Prpf31^+/-^* retinal samples.

### Prpf31 deficiency disturbs the mitochondrial respiratory chain

As mitochondrial SOD2 was the only detoxifying enzyme for which the expression levels were downregulated both at the mRNA and protein levels, we investigated if components of the mitochondrial respiratory chain were also affected. Usually, mitochondrial disorders are associated with mitochondrial respiratory chain impairment leading to ROS production, and can affect RPE and retinal cells (Brown et al., 2019; Decanini et al., 2007; Ferrington et al., 2017; He et al., 2010; Huang et al., 2023; Kowluru, 2013; Toms et al., 2019). Various subunits of the 5 respiratory chain complexes were analyzed: the NADH-ubiquinone oxidoreductase chain 4 (ND4) in complex I, the cytochrome C oxidase subunit 4 (CoxIV) in complex IV and the ATP synthase 6 (ATP6) from the ATP synthase complex V. *Nd4* showed some gene expression deregulation in *Prpf31^+/-^* retinal samples (Fig. 2 A). *CoxIV* was overexpressed in *Prpf31^+/-^* RPE/choroid fractions compared to wildtype at all ages tested, while *Atp*6 was overexpressed in *Prpf31^+/-^* RPE/choroid fractions at 3 months before being underexpressed at 6 and 12 months of age. Interestingly, and in contrast to gene expression, at the protein level COXIV was downregulated at 3, 6 and 12 months in *Prpf31^+/-^*RPE/choroid samples (Fig. 2 B). The decreased synthesis of ATP6 was confirmed at the protein level at 6 and 12 months of age. None of these 3 proteins showed any significant variation of their expression in corresponding retinal samples at all ages tested.

**Figure 2.**
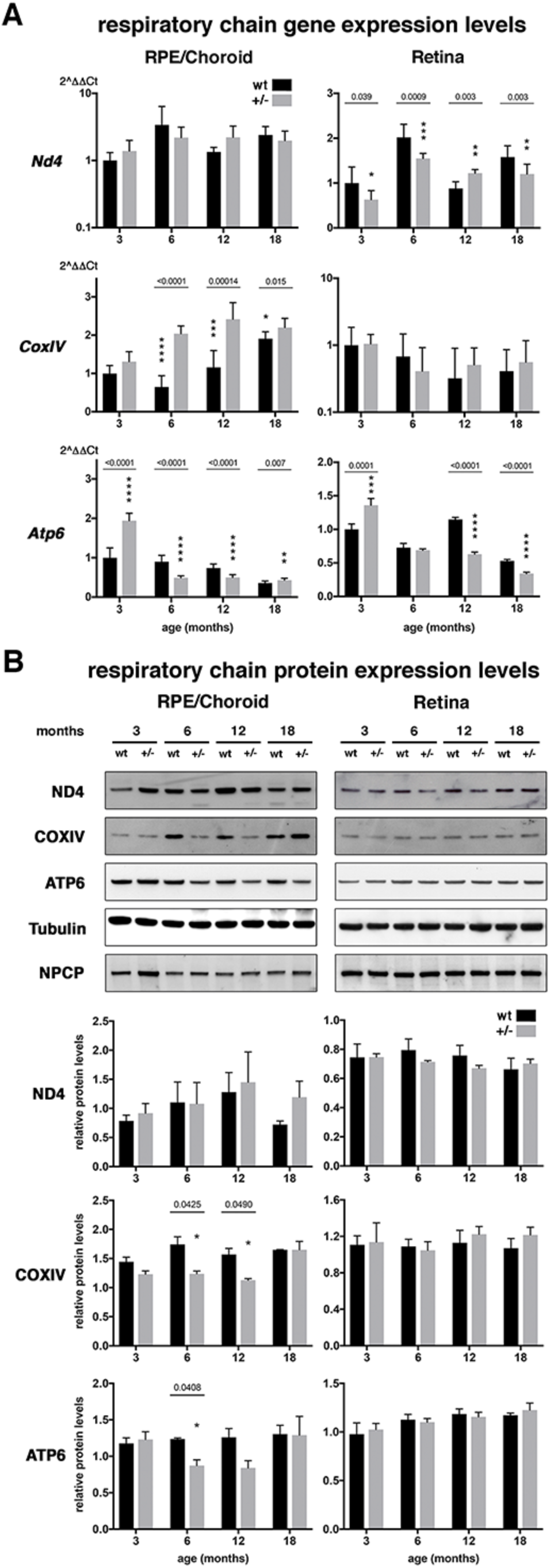
Alteration of the expression profiles of mitochondrial respiratory chain components in RPE/choroid and retina tissues from *Prpf31^+/-^* mice. RT-qPCR **(A)** and immunoblot **(B)** expression profiles of ND4, COXIV and ATP6 in wildtype (wt) and *Prpf31^+/-^*(+/-) RPE/choroid and retina fractions at different ages (months) as indicated. Tubulin and NPCP were used as internal loading controls for immunoblots. Data expressed as 2^ΔDCt relative to wt control samples at 3 months (A) or as relative protein levels compared to a control sample (B). Immunoblot band sizes (B): ND4 42-53kDa, COXIV 25kDa, ATP6 60kDA, tubulin 60kDa, NPCP 70kDa. Mean ± s.d., n = 5-12 (A) and n = 3-5 (B), multiple *t* test with significance threshold values set as * *P* < 0.05, ** *P* < 0.01, *** *P* < 0.001, **** *P* < 0.0001.

### Prpf31 deficiency affects several cellular energy sources

RPE and retinal cells are highly metabolic and require high amounts of energy to fulfill their different functions. The most important function of the mitochondria is the biosynthesis of adenosine triphosphate (ATP), even more in RPE cells for which glycolysis is not the main energy source (Kumagai et al., 1994; Swarup et al., 2019). Given the defects observed in components of the mitochondrial respiratory chain, we decided to analyze the mitochondrial activity in fresh *ex vivo* RPE/choroid punches using the Seahorse apparatus and Cell Mito Stress Test kit at 3, 6, 12 and 18 months. In this method, 3 drugs are successively injected –oligomycin, FCCP and rotenone/antimycin A– that inhibit the ATP synthase, the proton channel and complexes I/III, respectively. Amounts of oligomycin, FCCP and rotenone were titrated in preliminary experiments to determine optimal concentrations and measurement step timelines for our live tissues. Oxygen consumption rate (OCR) variations after the addition of each drug were measured which allowed the calculation of various parameters of the mitochondrial respiratory chain function. The extracellular acidification rate (ECAR) can also be calculated, giving information on extracellular CO_2_ and lactate accumulation. Overall, OCR and ECAR in *Prpf31^+/-^* RPE/choroid were lower than in wt controls (Fig. 3 A). Among all characteristics analyzed, 2 main changes were identified with a decrease in the basal respiration (BR, ∼45-64%) and in ATP production (ATPp, ∼53-64%), while other parameters such as maximal respiration, spare respiratory capacity, proton leak and non-mitochondrial oxygen consumption were globally not affected (Fig. 3 B).

**Figure 3.**
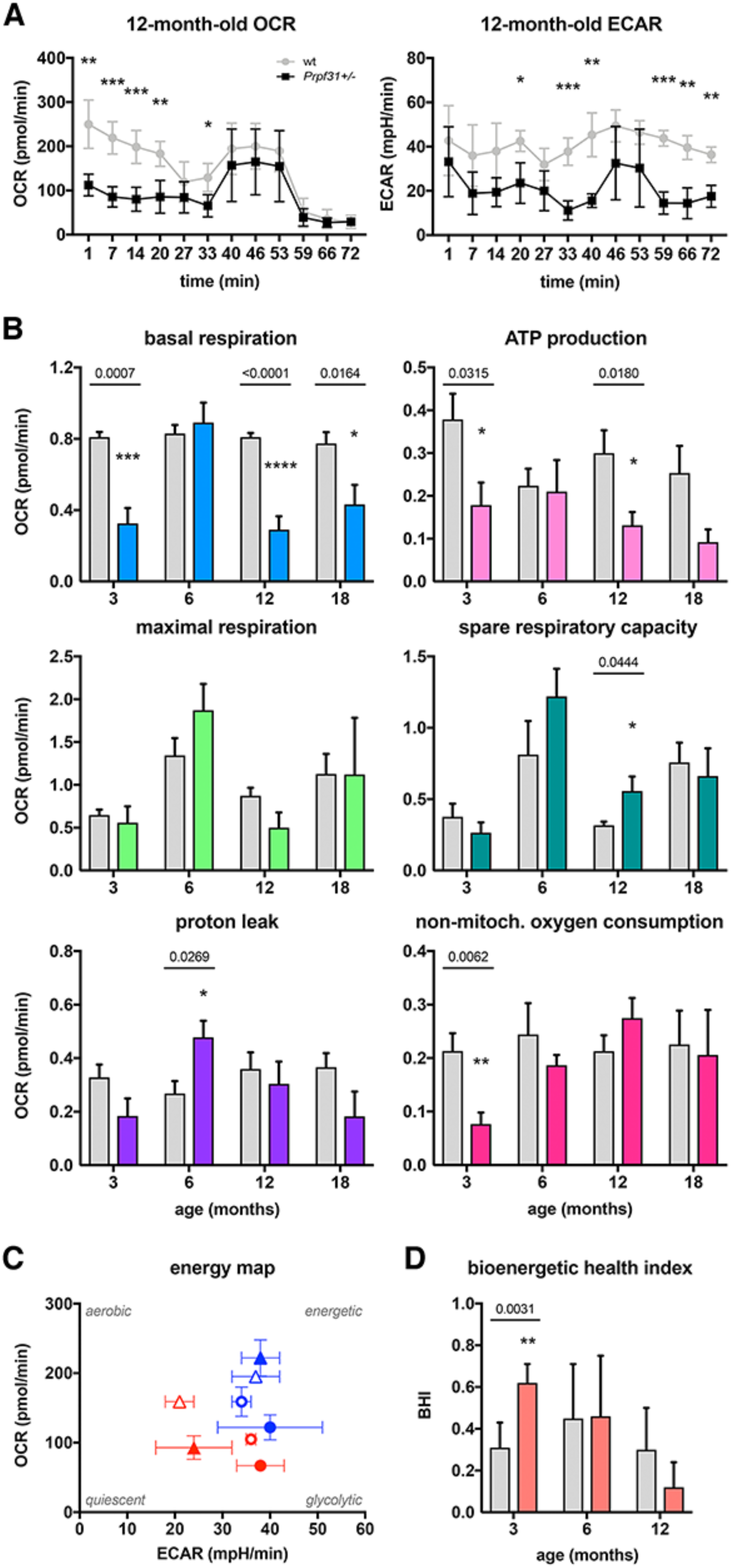
Decreased mitochondrial function and bioenergetic health index in RPE/choroid tissues from *Prpf31^+/-^* mice. **(A)** Representative OCR (left, pmol/min) and ECAR (right, mpH/min) profiles of 12-month-old wildtype (wt) control (grey line) and *Prpf31^+/-^*(black line) RPE/choroid punches. **(B)** Basal respiration (top left, blue), ATP production (top right, light pink), maximal respiration (middle left, light green), spare respiratory capacity (middle right, dark green), proton leak (bottom left, purple) and non mitochondrial oxygen consumption (bottom right, dark pink) profiles of 3-, 6-, 12- and 18-month-old wildtype control (grey bars) and *Prpf31^+/-^* (colored bars) RPE/choroid punches. All are expressed as OCR measures in pmol/min. **(C)** ECAR/OCR energy map of 3-(circles) and 12-month-old (triangles) wildtype control (blue) and *Prpf31^+/-^* (red) RPE/choroid samples under basic (full symbols) and stressed conditions (empty symbols). **(D)** Bioenergetic health index profile of 3-, 6- and 12-month-old wildtype control (grey bars) and *Prpf31^+/-^* (orange bars) RPE/choroid punches. Mean ± s.d., n = 4-7 (A,B,D), multiple *t* test with significance threshold values set as * *P* < 0.05, ** *P* < 0.01, *** *P* < 0.001, **** *P* < 0.0001.

Intriguingly, these changes in BR and ATPp were detected at 3, 12 and 18 months but not at 6 months, while the proton leak factor was increased at this exact age of 6 months. We thus extended our studies to decipher this discrepancy by producing the energy map and calculating the bioenergetic health index (BHI) (Fig. 3 C, D). Interestingly, these allowed us to show that *Prpf31^+/-^* RPE/choroid samples were evolving from a glycolytic (3 months) to a quiescent state (12 months), while wt control samples mostly remained midway between the energetic and glycolytic states using slightly less glycolysis with time (Fig. 3 C). In parallel, *Prpf31^+/-^*RPE/choroid samples BHI progressively decreased with age: higher than wt controls at 3 months, it was equivalent to wt at 6 months and then decreased at 12 months (Fig. 3 D).

These mitochondrial defects led us to examine other energy sources such as glucose and fatty acids, which could be used by the cells to bypass the deficiency in oxidative phosphorylation (OXPHOS). For that purpose, we used both the Glycolytic Rate Assay and Mito Fuel Flex Test Kits to assess these separate aspects of energy production. The glycolytic rate profile was obtained using the sequential injection of rotenone/antimycin A, blocking complexes I and III, and 2-deoxy-D-glucose (2-DG), blocking glycolysis. The glycolytic proton efflux rate (GlycoPER) is the rate of protons extruded into the extracellular medium during glycolysis, a combined measure of OCR and ECAR. Glycolysis analysis results followed the BHI trend, with a progressive decrease in both basal and compensatory glycolysis in *Prpf31^+/-^*RPE/choroid samples between the ages of 3 and 12 months, while wt controls remained stable with time (Fig. 4 A). This evolution was observed directly on the ECAR profiles at these different ages. Similarly to what we observed for OXPHOS, these different glycolysis data confirmed that at 6 months of age mutant and control samples did not differ. We next analyzed tissues requirement in fatty acids to meet the metabolic demand, known as fatty acid dependency. It was analyzed by adding etomoxir –an inhibitor of the long chain fatty acid oxidation–, followed by a combination of BPTES –inhibitor of the glutamine oxidation pathway– and UK5099 –inhibitor of the glucose oxidation pathway. Our results showed an overall decrease in fatty acids requirements for energy production in *Prpf31^+/-^* mice at all ages, and the most important diminition with target inhibitors at the age of 3 months (Fig. 4 B).

**Figure 4.**
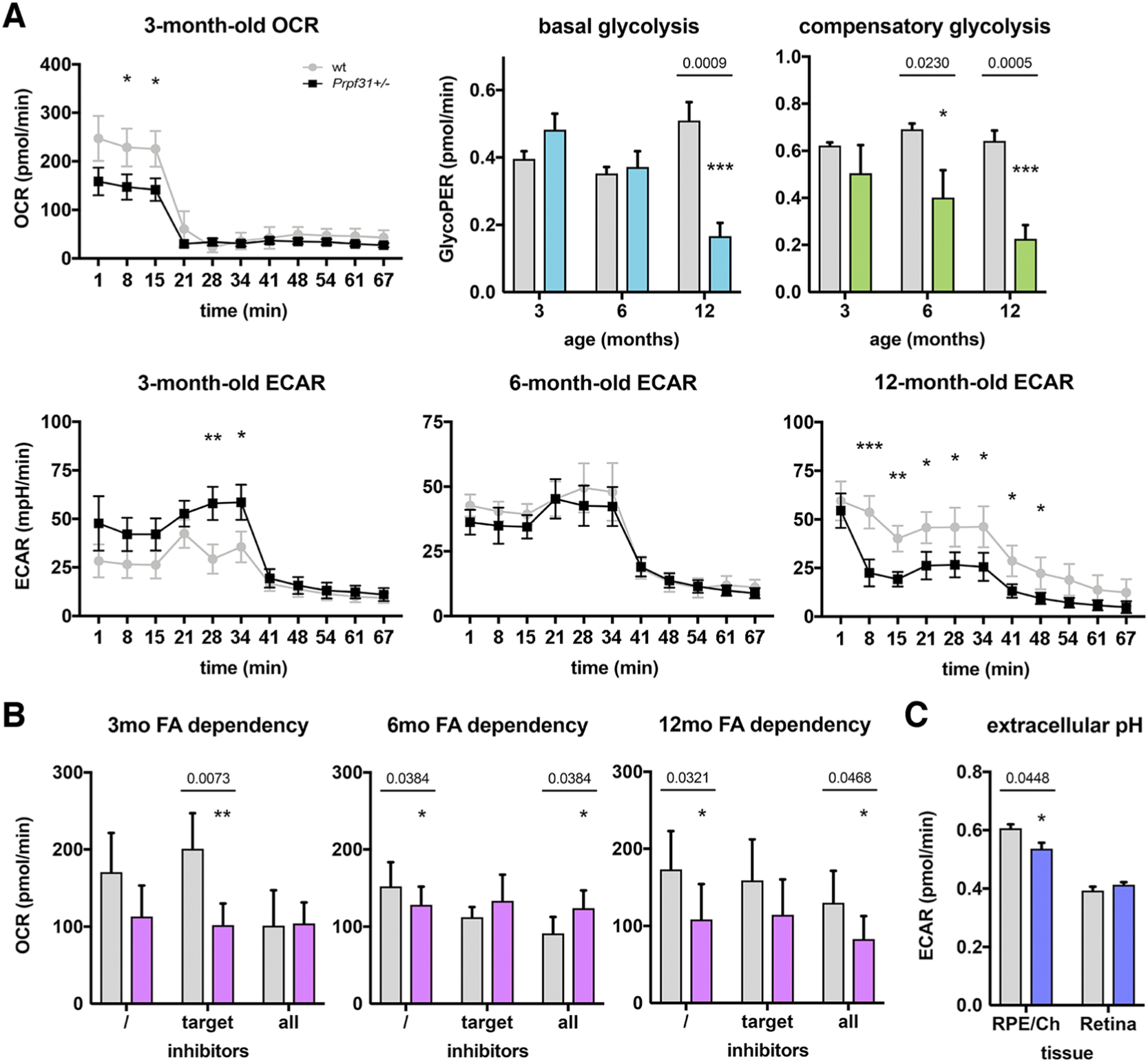
Decreased glycolysis and fatty acids dependency in RPE/choroid tissues from *Prpf31^+/-^*mice. **(A)** Representative OCR (top, pmol/min) and ECAR (bottom, mpH/min) profiles of 3-, 6- and 12-month-old wildtype (wt) control (grey line) and *Prpf31^+/-^* (black line) RPE/choroid punches. Basal (top left, blue) and compensatory (top right, green) glycolysis profiles of 3-, 6- and 12-month-old wildtype control (grey bars) and *Prpf31^+/-^* (colored bars) RPE/choroid punches expressed as GlycoPER measures in pmol/min. **(B)** Fatty acids dependency profiles of 3- (left), 6- (middle) and 12-month-old (right) wildtype control (grey bars) and *Prpf31^+/-^* (purple bars) RPE/choroid punches expressed as OCR measures in pmol/min. Inhibitors used: / = none, target = etomoxir followed by BPTES/UK5099, and all = all 3 inhibitors simultaneously. **(C)** Extracellular pH profile of wildtype control (grey bars) and *Prpf31^+/-^* (blue bars) RPE/choroid (RPE/Ch) and retina samples expressed as ECAR measures in mpH/min. Mean ± s.d., n = 4-7 (A,B) and n = 3 (C), multiple *t* test with significance threshold values set as * *P* < 0.05, ** *P* < 0.01, *** *P* < 0.001, **** *P* < 0.0001.

RPE and retinal cells are highly dynamic, hence our protocol required 1-mm punches to maintain enough oxygen before the addition of drugs to allow measures of differences during the whole assays. Therefore, this size of sample did not allow the analysis of retinal tissues too fragile to be cut and conserved in a state satisfying enough to conduct these very sensitive tests. However, and in order to validate if these changes were restricted to RPE/choroid samples, we utilized another kit (MitoXpress Xtra Oxygen Consumption Assay) using multiplates compatible with the use of full RPE/choroid cups and retinas. This allowed us to show that while the extracellular pH was indeed decreased in *Prpf31^+/-^* RPE/choroid samples, it was not the case in corresponding separated retinal samples (Fig. 4 C).

As we showed previously that the phagocytic defect present in RPE cells did not affect peritoneal macrophages which use the same molecular machinery for phagocytosis of apoptotic cells (Farkas et al., 2014), we asked wether these metabolic dysfunctions were present in peritoneal macrophages. Interestingly, neither the mitochondrial function, glycolysis nor fatty acids usage were affected in *Prpf31^+/-^* peritoneal macrophages (Fig. S1). The overall mitochondrial activity measured using MitoTracker probes was also similar between mutant and control macrophages.

To further investigate the energetic state of *Prpf31^+/-^*mice at the whole body level, mice were independantly studied for 1 week in metabolic cages and various parameters monitored. Our results confirmed that at the global body level mutant and control mice had the same body weights, food consumption levels, respiratory exchange rates and energetic expenses, both during the day and at night (Fig. S2).

### Prpf31 deficiency induces endoplasmic reticulum stress and activation of the unfolded protein response

In the absence of an efficient antioxidant system and with the concommitant dysfunction of mitochondrias, oxidative stress contributes to the accumulation of abnormal misfolded proteins in the ER, leading to ER stress (Malhotra and Kaufman, 2007). Deregulation of ER homeostasis activates the UPR by the sequential mobilization of 3 pathways leading to different outcomes. Normally, proper protein folding in ensured by chaperonnes of the Hsp family or the immunoglobulin heavy chain binding protein/glucose-regulated protein 78 (Bip/Grp78), which is bound to 3 main stress transducer proteins anchored at the ER membrane: inositol requiring enzyme-1 alpha (IRE1α), protein kinase R-like endoplasmic reticulum kinase (PERK) and activating transcription factor 6 (ATF6) (Zhang et al., 2014; Ong and Logue, 2023). Upon the detection of unfolded or misfolded proteins, Bip/Grp78 detaches from these 3 stress sensors, thus activating their downstream pathways –Xbp1 for IRE1α, Atf4 for PERK and the cleavage of Atf6 in the Golgi apparatus–, ultimately leading to the transcription of UPR target genes in the nucleus.

To assess the presence of ER stress we thus investigated some components of the UPR implicated in this process at the mRNA and protein levels. We detected a deregulation of the *Bip* gene in RPE/choroid as well as retinal tissues at 3, 6 and 12 months of age (Fig. 5 A). Expression of *Atf4* was mostly similar between control and mutant samples, except for retinal samples at 6 months of age. The gene expression profile of the *C/EBP-homologous protein* (*Chop*) –the main downstream effector for Atf4– followed the same trend than Atf4 in wt RPE/choroid fractions, decreasing between the ages od 3 and 12 months before increasing again at 18 months. *Chop* was slightly overexpressed at 3 months in *Prpf31^+/-^* RPE/choroid fractions, and markedly in retinal samples at 6 months and slightly at 12 months. *Atf6* expression appeared to vary with age in wt RPE/choroid control samples, decreasing between 3 and 12 months and increasing back at 18 months. Differences were observed in *Prpf31^+/-^*RPE/choroid samples at 6 months, and in retinal samples at all ages, first decreased at 3 months, then increased at 6, 12 and 18 months.

**Figure 5.**
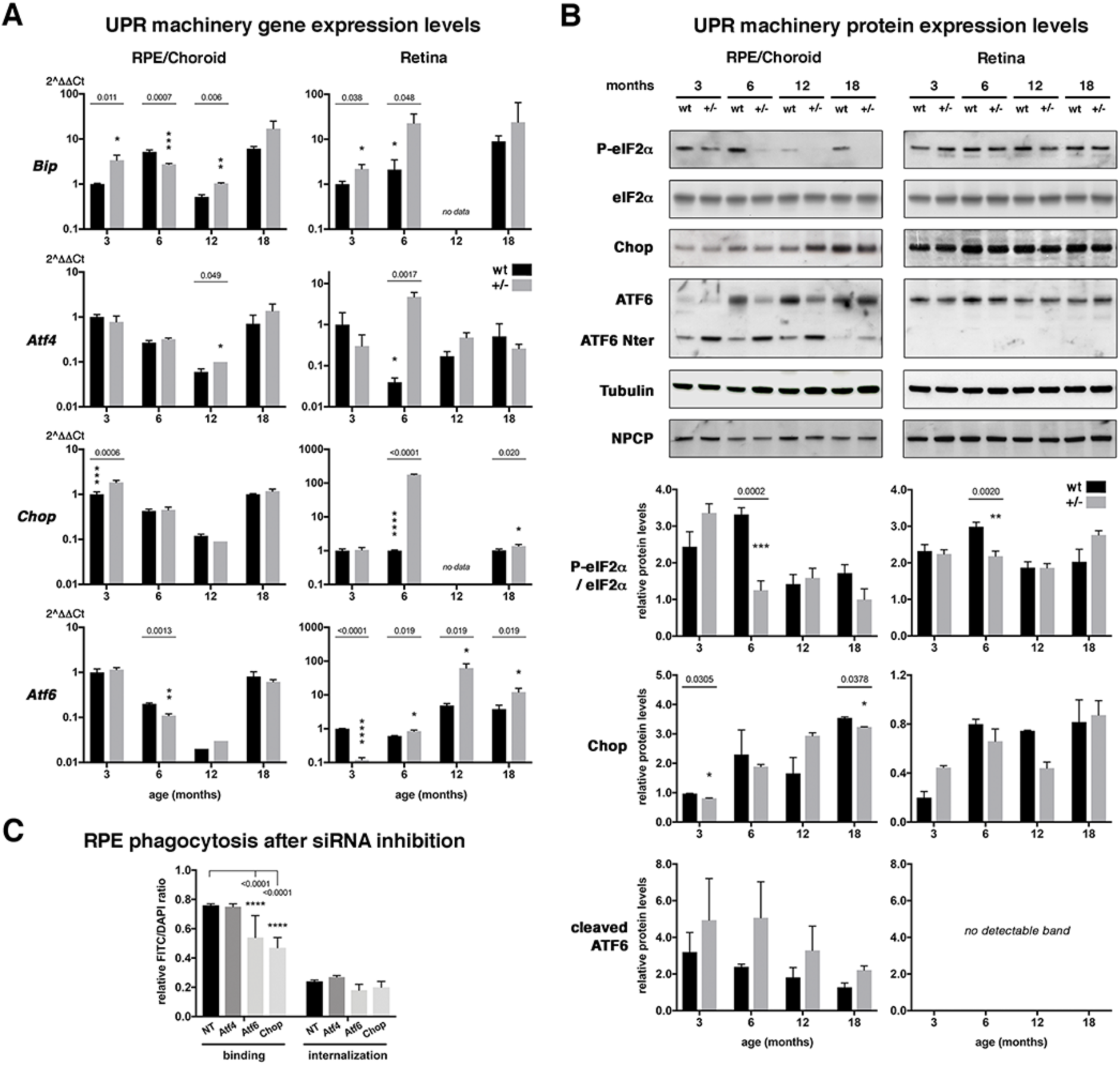
Activation of the unfolded protein response in RPE/choroid and retina tissues from *Prpf31^+/-^* mice and impact of ER stress on RPE phagocytosis. RT-qPCR **(A)** and immunoblot **(B)** expression profiles of BIP, ATF4, CHOP, ATF6, eIF2α and phospho-eIF2α in wildtype (wt) and *Prpf31^+/-^* (+/-) RPE/choroid and retina fractions at different ages (months) as indicated. Tubulin and NPCP were used as internal loading controls for immunoblots. Data expressed as 2^ΔΔCt relative to wt control samples at 3 months (A) or as relative protein levels compared to a control sample (B). Immunoblot band sizes (B): eIF2α/P-eIF2α 40kDa, ATF6 70kDa, cleaved ATF6 40kDA, Chop 28kDa, tubulin 60kDa, NPCP 70kDa. **(C)** RPE binding and internalization phagocytosis profile of RPE-J cells after siRNA downregulation of *Atf4*, *Atf6* and *Chop* genes when compared to a control siRNA (Non-targeting, NT) as indicated. All phagocytosis results are expressed as relative FITC/DAPI ratios. Mean ± s.d., n = 3-5 (A, except n = 2 in *Prpf31^+/-^* RPE/choroids for *Atf6* at 6 and 12 months, for Chop at 12 months, and in *Prpf31^+/-^* retinas for *Atf6* at 3 and 6 months), n = 2-4 (B), and n = 4-5 (C), multiple *t* test with significance threshold values set as * *P* < 0.05, ** *P* < 0.01, *** *P* < 0.001, **** *P* < 0.0001.

Interestingly, when studying the related proteins more information can be obtained than the mere expression levels. Indeed, some components of the UPR can be phosphorylated such as eIF2α, thus activating the ATF4 pathway, or cleaved, such as ATF6 showing its direct activation. In our samples, eIF2α phosphorylation appeared to decrease with age in wt RPE/choroid tissues, while being lower at all ages in *Prpf31^+/-^* and similar and stable in all retinal tissues (Fig. 5 B). We detected a progressive increase in Chop proteins expression with time in all samples, with a more variable profile in *Prpf31^+/-^* RPE/choroid samples. The ATF6 protein profile is interesting because ATF6 was more cleaved in *Prpf31^+/-^* RPE/choroid samples, essentially between the ages of 3 and 12 months, while no cleavage at all could be detected in corresponding retinal samples.

We then asked the question whether proteasome deficiencies would directly impact the phagocytic function of RPE cells. For that purpose, we analyzed the capacity of RPE-J cells treated with *Atf4*, *Atf6* and *Chop*-specific siRNAs to tether and ingest POS *in vitro*. *Atf6* and *Chop* downregulation significantly affected POS binding by RPE cells, while internalization was also decreased but non significantly (Fig. 5 C).

### Prpf31 deficiency sensitizes RPE cells to oxidative stress

We previously detected a ∼40% decrease of POS phagocytosis by *Prpf31^+/-^*RPE cells in primary cultures, as well as for the human ARPE-19 cell line down-regulated for *PRPF31* and in iPSC-RPE cells derived from patients (Farkas et al., 2014; Rodrigues et al., 2022). Hence, we wondered if ER stress had a negative effect that could worsen the impact of *Prpf31* knockdown on the phagocytic process. We used a common ER stressor, tunicamycin, in concentrations that affect neither phagocytosis nor cell viability in basal conditions, i.e. in presence of normal amounts of Prpf31. As demonstrated previously, RPE-J cells treated with *Prpf31*-targeting shRNAs expressed less Prpf31 proteins and displayed a decreased phagocytic capacity (≈40-50%) both for the binding and the internalization steps (Fig. 6) (Farkas et al., 2014). Interestingly, non-offensive doses of tunicamycin appeared to affect POS internalization and aggravated the loss of POS uptake when Prpf31 was also downregulated.

**Figure 6.**
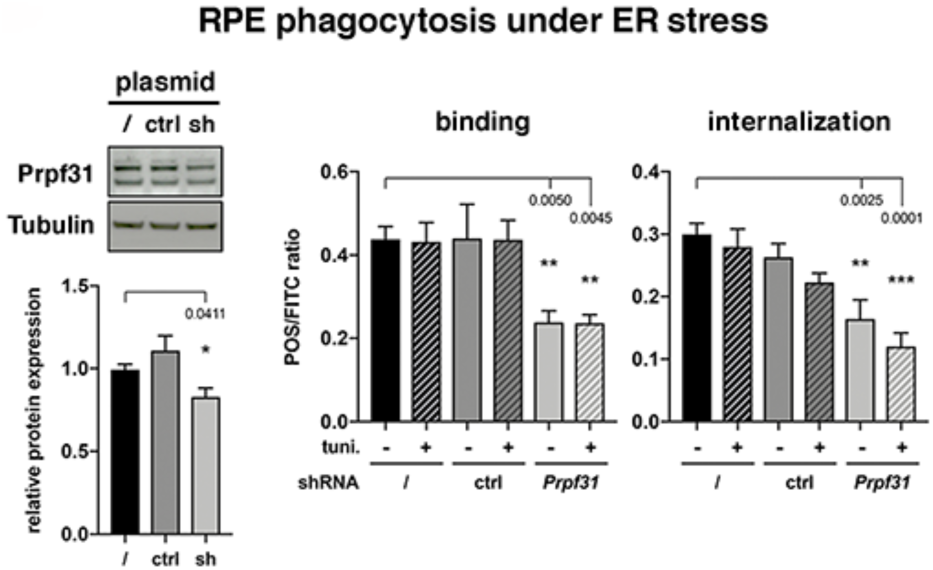
Impact of ER stress on RPE phagocytosis. Immunoblot downregulation validation and binding and internalization phagocytic profiles of RPE-J cells treated with control (ctrl) or *Prpf31*-specific shRNAs when compared to untreated cells (/) with (+) or without (-) the addition of ER stressor tunicamycin as indicated. All phagocytosis results are expressed as relative FITC/DAPI ratios. Mean ± s.d., n = 3-5, multiple *t* test with significance threshold values set as * *P* < 0.05, ** *P* < 0.01, *** *P* < 0.001, **** *P* < 0.0001.

## Discussion

PRPF splicing factors constitutes a family of ubiquitously expressed proteins involved in the formation of stable U4/U6.U5 tri-snRNPs and the spliceosomal B complex required for spliceosome activation. Surprisingly, *PRPF* mutations result solely in retinal-specific phenotypes. To this day, despite several studies there are no definitive evidence that define the primary affected cell type in the retina between RPE and photoreceptors, or provide clear insights into the precise pathomechanisms of this disease (Azizzadeh Pormehr et al., 2020; Farkas et al., 2014; Graziotto et al., 2011; Hebbar et al., 2021; Huranová et al., 2009; Maxwell et al., 2021; Mordes et al., 2007; Rodrigues et al., 2022; Valdés-Sanchez et al., 2019; Wheway et al., 2015; Wilkie et al., 2006; Yin et al., 2011).

In the *Prpf31^+/-^* mouse model, RPE phagocytosis and its circadian rhythm are disturbed since an early age, and later on RPE cells show signs of cellular stress (Graziotto et al., 2011; Farkas et al., 2014). Despite the difference in the timing of pathological development between Human patients and mice, similar mechanisms such as phagocytosis are affected in both species showing the *Prpf31^+/-^* model can be used as a paradigm and could be ideal to decipher the precise chronology of pathological development of *PRPF*-related adRP (Farkas et al., 2014). Hence, in this study, we analyzed further the *Prpf31^+/-^* mouse model focusing on several stress pathways usually present in cells.

In RPE cells, OXPHOS is the major source of energy as the majority of the glucose coming from choroidal vessels is translocated to the retina to be redistributed to RPE cells as pyruvate (Adijanto and Philp, 2014; Kumagai et al., 1994; Léveillard et al., 2019; Swarup et al., 2019). *Prpf31*^+/-^ RPE/choroid tissues show diminished mitochondrial SOD2, COXIV and ATP6 expression levels. These are correlated with impaired mitochondrial function, mainly through decreased respiration and ATP production as early as 3 months of age. *Sod2^-/-^* mice have previously been shown to display oxidative stress, mitochondrial dysfunction in both RPE and photoreceptors, as well as mitochondria and RPE morphology alterations (Brown et al., 2019). In this model, RPE cells undergo a metabolic switch from OXPHOS to glycolysis as a compensatory mechanism for ATP production. However, in our model glycolysis diminishes with time, as well as the use of fatty acids for energy production. This is in accordance with the analysis of the BHI and energy map which show that *Prpf31*^+/-^ RPE/choroid tissues transition from a glycolytic to a more quiescent state while wt tissues stay midway between the glycolytic and the energetic states. Our results suggest that *Prpf31^+/-^* RPE/choroid samples undergo this metabolic switch around the age of 6 months, an age at which the function of mitochondria is similar between mutant and wt samples. Interestingly, comparison of mitochondrial activity with different tissues show a more pronounced and earlier phenotype in RPE cells than in the retina. In addition, peritoneal macrophage and overall body metabolisms were not affected, hereby validating further the tissue-specificity of the functional defect already observed for phagocytosis (Farkas et al., 2014).

Overall, *Prpf31^+/-^* RPE/choroid tissues display a general defect in energy production that might impact further several crucial energy-consuming cellular functions such as phagocytosis. Knowing the importance of mitochondria for ciliogenesis (Burkhalter et al., 2019), these results might also explain the limited ciliogenesis observed in iPSC-RPE cells derived from *PRPF31*-RP11 patients (Buskin et al., 2018; Maxwell et al., 2021; Wheway et al., 2015). Metabolic uncoupling –i.e. the reduced efficiency of OXPHOS to produce ATP– altering bioenergetics has previously been shown to take place in aged RPE cells *in vitro* as well as in diseased retinas (Ferrington et al., 2017; Fisher and Ferrington, 2018; He et al., 2010; Schütt et al., 2012), and to correlate with increased susceptibility to oxidative stress (Rohrer et al., 2016). Notably, mitochondria are a major source of free radicals through the production of ROS during the OXPHOS reaction, showing that energy production also contributes to oxidative stress (Turrens, 2003). Concomitantly, under oxidative stress conditions when detoxifying enzymes fail to neutralize ROS, cells block the OXPHOS and switch to glycolysis which is energetically more favorable (Anastasiou et al., 2011; Brand and Hermfisse, 1997; Molavian et al., 2016). In *Prpf31^+/-^* RPE/choroid we do observe a transition from OXPHOS to glycolysis, but ROS and oxidations still increase with especially on lipids, suggesting that other oxidative processes take place in these tissues.

In this context, we do show the induction of generalized oxidative stress with age in *Prpf31*^+/-^ mice, affecting proteins in RPE cells and lipids in the whole retina in aged animals. It has been shown previously that despite the fact that ROS can be short-lived in cells, their damages on cellular components, such as 4-HNE we observed in *Prpf31^+/-^* RPE and retinas, can be long-lived and a source of chronic stress (Handa, 2012). The increase in SOD2 expression at 3 months might suggest the activation of the oxidative stress response in mitochondria, while the downregulation of its expression from 6 months of age onwards suggest a disruption of the anti-oxidant response leading to oxidative stress. Indeed, cumulative effects of the imbalance in detoxifying enzymes expression such as SOD2, ER stress and UPR activation might aggravate the situation.

Defective mitochondrial function is often associated with ER stress, a normal process observed in aging cells (Ong and Logue, 2023). The ER is a central organelle involved in protein synthesis and maturation. Unfolded proteins accumulate in the ER and disturb its homeostasis, which may lead to cell death if the UPR fails to prevent the accumulation of misfolded proteins, including in RPE cells (Breckenridge et al., 2003; Kaarniranta et al., 2010). The UPR acts by delaying protein translation, degrading misfolded proteins, and increasing the production of chaperones involved in protein folding (Ron and Walter, 2007). *Prpf31*^+/-^ mice present oxidation of proteins observed via nitrotyrosine labeling– mostly in RPE cells, which might cause their subsequent aggregation in the ER (Malhotra and Kaufman, 2007; Malhotra et al., 2008). To confirm this hypothesis we analyzed the activation of the 3 UPR pathways via the expression/activation of different UPR sensors and effectors. Among these, normal aging activated the Atf4 pathway leading to Chop overexpression similarly between wt and mutant RPE/choroid tissues, potentially promoting autophagy, amino acid metabolism and anti-oxidant responses (Ong and Logue, 2023). Interestingly, activation of ATF6, which takes place by its cleavage in the Golgi apparatus, was marked in *Prpf31^+/-^*RPE/choroid but not retinal samples, suggesting the intervention of chaperones increasing protein folding capacity and the activation of anti-oxidant response genes (Ong and Logue, 2023). Notably, expression of the Hspa4l chaperone is upregulated and its association with mutant Prpf31 proteins was augmented in *Prpf31^A216P/+^* RPE cells (Valdés-Sanchez et al., 2019). The activation of this survival pathway linked to Atf6 might thus explain, at least in part, the absence of cell death in this heterozygous mouse model.

Protein synthesis is highly related to intracellular ATP levels, and it has been shown that a 15% lower level of ATP production leads to a 50% lower level of protein synthesis (Gronostajski et al., 1985; Wieser and Krumschnabel, 2001). In these conditions, the decrease in ATP levels we observed might correlate with the activation of the UPR in *Prpf31^+/-^* RPE/choroid cells, which has been shown to contribute to the reduction of the amounts of newly produced proteins and their translocation into the ER lumen (Kaufman et al., 2002). Moreover, the deregulation in *Bip/Grp78* gene expression, a glucose-regulated protein, might also be linked to the change in energetic metabolism and decrease in glycolysis. Furthermore, impairment of phagocytosis and mitochondrial function can be accompanied by dysregulation of cell proteostasis and redox homeostasis, as well as POS generation and accumulation of protein aggregates and lipids (Kaarniranta et al., 2020; Markitantova and Simirskii, 2025). This long-term modifications contribute to generalized oxidative stress and further RPE dysfunction by the slow down of multiple cell functions.

To address the impact of ER stress on phagocytosis we used sub-lethal doses of the ER stress inducer tunicamycin. We confirmed that stressing the ER did worsen the phagocytic defect of *Prpf31^+/-^* RPE cells. These results show the importance of ER homeostasis for the phagocytic activity of RPE cells. Phagocytosis *per se* does produces ROS in RPE cells or macrophages, and predominently upregulates catalase but not SOD activity (Miceli et al., 1994). The deregulated phagocytic rhythm in *Prpf31*^+/-^ RPE cells detected in very young animals (Farkas et al., 2014) might worsen oxidative stress production as it is the case in other models and systems both *in vivo* and *in vitro* (Dorey et al., 1989; Nandrot et al., 2004). As shown previously, oxidative and ER stress might in turn impact further the efficiency of phagocytosis, thus reinforcing oxidative stress levels as was observed in our model (Kaemmerer et al., 2007; Sun et al., 2006). Taken together, our results suggest the presence of a vicious circle of down-regulated functions, like phagocytosis and the energetic metabolism, leading to cellular and ER stress further worsening various cell activities leading to even more stress. In addition, our data suggest that energetic failure, oxidative imbalance and associated stress take place first and foremost in *Prpf31*^+/-^ RPE cells, thus reinforcing the idea that the RPE might be the primary cell type affected in splicing factor related-RP. Further studies characterizing specific transcripts modified in these cells could help characterizing the exact molecular mechanisms underlying these defects (Buskin et al., 2018; Mordes et al., 2007; Rodrigues et al., 2022; Tanackovic et al., 2011; Yuan et al., 2005). The “slow phenotype” of our heterozygous mouse model –devoid of any retinal degeneration unlike Human patients– constitutes an advantage and allows us to dissect the full series of cellular events, such as mitochondrial dysfunction and UPR activation, that will now have to be validated in Human iPSC-RPE cells. These studies could have a broader impact as more spliceosome-related genes are being identified in RP (Quinodoz et al., 2025; Ruzickova and Stanek, 2016), potentially leading to similar molecular defects as it has already been show for *PRPF3*, *6*, *8* and *31* (Farkas et al., 2014; Liang et al. 2022; Tanackovic et al., 2011).

## Methods

### Animals and Tissue Collection

Animals were handled according to the Association for Research in Vision and Ophthalmology Statement for the Use of Animals in Ophthalmic and Vision Research. Protocols were approved by the Charles Darwin Animal Experimentation Ethics Committee from Sorbonne Université and the French Ministry for Education, Higher Studies and Research (APAFIS#1631-2015090415466433 and APAFIS#20191-2019040311402311). For experiments, wildtype and *Prpf31^+/-^*mice aged from 3 to 18 months were sacrificed by CO_2_ asphyxiation. For immunohistochemistry experiments, carefully dissected eyes were fixed in Davidson fixative for 1hr at 4°C, the cornea opened, eyes fixed for 3 more hours at 4°C, lens and cornea removed and then eyecups further fixed for 3hrs at 4°C. After overnight dehydration steps on the Spin Tissue Processor STP 120 (Myr, Thermo Scientific), eyecups were embedded in paraffin and 5-μm sections were cut. For gene and protein expression experiments, eyes were collected and swiftly dissected in HBSS without CaCl_2_ and MgCl_2_. After removal of the lens, the retina was carefully separated from the rest of the cup (RPE/choroid) and both parts were frozen separately in liquid nitrogen. One eye from each animal was used for gene expression levels assessment and the fellow eye for protein levels analysis (see respective sections below). For metabolic flux analysis, surrounding tissues were cleaned out and either 1-mm RPE/choroid punches were dissected out and immediately transferred to the assay cartridge while letfover tissues flash frozen in liquid nitrogen, or full RPE/choroid and separated retina were transferred into individual wells of a 96-well plate for assay processing (see corresponding methods section).

### Immunohistochemistry

After removing the paraffin in 2 baths each of Ottix Plus and Ottix Shaper (Diapath), slides were rehydrated and treated with boiling 1X citrate buffer for 20min. Membranes were permeabilized in 0.3% Triton X-100 in 1X TBS for 5min followed by 2 5-min washes in 1X TBS. The blocking step of non-specific signals was performed with 4% bovine serum albumin in 1X TBS prior to the overnight incubation with primary antibodies at 4°C (Supplementary Table 1). Alexa-conjugated secondary antibodies (488 and 594, Molecular Probes, Life Technologies) donkey anti-rabbit and donkey anti-mouse (all 1:300) were used to amplify signals. Nuclei were stained with DAPI (1mg/mL in 1X PBS) and slides were mounted using Vectashield (Vector Laboratories). Images were acquired with an upright Olympus FV1000 laser-scanning confocal microscope equipped with standard PMTs and high-sensitivity GaAsP detectors via the Fluoview 2.1c software, using a 2X zoom on a PLAPON 60X super corrected objective at room temperature.

### RNA Extraction, Reverse Transcription and Real-time Quantitative PCR

Total RNAs from RPE/choroid and retina fractions were isolated using the Illustra RNAspin Mini kit (GE Healthcare) according to the manufacturer’s instructions and using a second DNAse step. 500ng of RNAs were reverse-transcribed in the presence of oligo(dT) primers in a final volume of 50µL using the Reverse Transcription System (Promega) as instructed. Relative expression of the various genes was quantified by real-time PCR using the SYBR Green PCR Master Mix (Applied Biosystems) and 100nM of the appropriate primers (Supplementary Table 2) on a StepOne Plus apparatus (Applied Biosystems). The *ribosomal protein Rho0* (*Rplp0*) housekeeping gene was used as an internal control for normalization. Relative amounts of each gene were calculated using the 2^-ΔΔCt^ method and expression levels for wildtype mice at 3 months were set to 1 and used as reference as specified in each figure legend.

### Protein Extraction and Immunoblot Analysis

RPE/choroid and retina samples were homogenized on a shaker at 1500rpm at 4°C for 30min in a lysis buffer composed of 50mM Tris/HCl pH 7.8, 150mM NaCl, 2mM CaCl_2_, 1mM MgCl_2_, 1% Nonidet P40 and 1% Triton X-100, supplemented with a protease inhibitor cocktail (Sigma) and 1% PMSF. Lysates were cleared by centrifugation at 14,000g for 10min at 4°C. Proteins were quantified using the Bradford method (Sigma) and a BSA range, 12µg of proteins separated on 4-12% NuPAGE Bis-Tris protein gels (Invitrogen), and transferred onto nitrocellulose membranes (Amersham). Membranes were blocked for 1hr at room temperature in 10% non-fat dry milk in 1X TBS, and subsequently incubated overnight at 4°C with primary antibodies (Supplementary Table 1). Then, blots were incubated for 1hr with the appropriate secondary antibody conjugated to the horseradish peroxidase (1:2,000). Immunoreactive bands were visualized using the Western Lightning Plus-ECL chemiluminescence detection system (Perkin Elmer) and Amersham Hyperfilm ECL films (GE Healthcare). Quantitative analysis of detected bands was performed using ImageJ. Duplicates of an identical control sample were loaded on each immunoblot and used as reference for protein quantifications and comparisons between different sample series.

### Real-Time Cell Metabolic Analysis and *In Vivo* Metabolic Phenotyping

Eyes were collected from wildtype and *Prpf31^+/-^* mice at different ages and RPE/choroid and retina fractions separated as described in the Tissue Collection section. For each animal, freshly dissected mm RPE/choroid punches (3 total for 2 eyes) were collected in individual wells of a FluxPak Seahorse XFp cartridge in presence of Seahorse XF base medium minimal DMEM without phenol red (Agilent Technologies) supplemented with 1mM pyruvate, 2mM glutamine and 10mM glucose, and adjusted to pH7.4. Before each assay, punches were equilibrated at 37°C in a non-CO_2_ incubator for 1hr. For the XFp Cell Mito Stress Test Kit (Agilent Technologies), we used the mitochondrial inhibitors oligomycin (1.5μM), FCCP (carbonyl cyanide 4-(trifluoromethoxy)-phenylhydrazone, 0.5µM) and rotenone/antimycin A (0.5µM) (Enderlin et al., 2024). Oxygen Consumption Rate (OCR) and ExtraCellular Acidification Rate (ECAR) were automatically calculated and recorded by the Seahorse Report Generator online software (Agilent Technologies), and expressed as pmol/min and mpH/min, respectively. The original program was modified to meet our experiment requirements: each experiment was separated into 4 intervals between drugs addition, and 3 measurements were performed for each step and each cycle of 3min mixing, 1min waiting and 2min measuring. The energy map was graphically generated by plotting ECAR (x-axis) against OCR (y-axis) values in basic (mean of the first 3 measures) and stressed (mean of the 3 values after addition of oligomycin and FCCP, respectively) conditions for each genotype at 3 and 12 months of age. The bioenergetic health index (BHI) was calculated using values obtained from the XFp Cell Mito Stress Test Kit using the following formula: BHI = (ATP production x spare respiratory capacity)/(proton leak × non-mitochondrial oxygen consumption). The XFp Glycolytic Rate Assay (0.5µM rotenone/antimycin A, followed by 50µM 2-deoxy-D-glucose) and XFp Mito Fuel Flex Test (fatty acids dependency: 4µM etomoxir, followed by 3µM BPTES and 2µM UK5099 inhibitors of the 2 alternative pathways) Kits (Agilent Technologies) were used according to the manufacturer’s instructions. Glycolysis analysis results are expressed as glycolytic proton efflux rate (GlycoPER) in pmol/min and fatty acids dependency as OCR in pmol/min.

Oxygen Consumption Assays on full RPE/choroid and retina fractions were conducted in 100µL final complete seahorse media with 10µL of reconstituted MitoXpress Xtra – Oxygen Consumption Assay reagent (Agilent Technologies) in a 96-well microplate following the provided protocol. Individual experimental wells were sealed by adding 2 drops of pre-warmed mineral oil and the plate was monitored on a microplate reader (Infinite M1000, Tecan) using the TR-F mode at 37°C (380±9nm excitation and 650±20nm emission filters). Signals from each well were measured repetitively every 2 minutes over 90min with 30µs of lag time and 100µs of integration time.

MitoTracker Red FM (active mitochondria) and MitoTracker Green FM (all mitochondria) probes (Molecular Probes, Invitrogen) were resuspended in DMSO at a concentration of 1mM, and were diluted sequentially at 1µM and 200nM in Seahorse XF base medium. 90µL of each 200-nM probe were loaded per well, and tissues/cells were incubated in the dark for 30min at 37°C. After 2 washes with Seahorse XF base medium, fluorescence signals were measured on a microplate reader (Infinite M1000, Tecan) using the fluorescence mode at 37°C (581±5nm or 490±5nm excitation and 640±5nm or 516±5nm emission filters for MitoTracker Red FM and MitoTracker Green FM, respectively).

After all assays, protein levels were determined for each sample to normalize the results. A second normalization step was performed for each analysis against the average of the wildtype sample starting values.

For the monitoring of full body metabolism mice were followed-up individually on a Phenomaster apparatus (TSE Systems). Mice (2 males and 4 females from each genotype) were first habituated with the drinking probe in regular cages for 3 days and then with the calorimetric cages for 3 more days individually. Mice were monitored for 5 days for the following data: direct calorimetry measuring the volumes of O_2_ consumed CO_2_ produced (respiratory exchange rate = vCO_2_/vO_2_), water and food consumption, spontaneous vertical and horizontal locomotion. At the beginning and the end of the experiment, mouse lean/fat mass and composition were evaluated by nuclear magnetic resonance.

### Cell Culture and Transfection

The rat RPE-J cell line (ATCC) was maintained in DMEM (Gibco) supplemented with 4% CELLect Gold FCS (MP Biomedicals), 10mM HEPES, 1% non-essential amino acids and antibiotics (100U/mL penicillin and 100μg/mL streptomycin) (all from Gibco) in a humidified atmosphere containing 5% CO_2_ at 32°C. For transfection and phagocytosis experiments, RPE cells were plated on Alcian blue-coated (Sigma) 96-well plates. Cells were transfected twice 24 and 72 hours post splitting with validated ON-TARGETplus SMARTpool siRNAs (Atf4 L-099212-02, Atf6 L-083071-02, Chop L-088282-02, Non-Targeting control D-001810-10) with the DharmaFECT 4 siRNA Transfection Reagent (Dharmacon, Horizon Discovery), and phagocytosis assays performed 24 hours after the second transfection as described previously (Parinot et al., 2024). Alternatively, cells were transfected 24 hours post splitting with the different expression plasmids coding for *Prpf31*-specific (TGAAGATTGAGGAGTATATTCTCTTGAAATATACTCCTCAATCTTCA) and luciferase control (CTTACGCTGAGTACTTCGATTCAAGAGATCGAAGTACTCAGCGTAAG) shRNAs for 5hrs using Lipofectamine 2000 (Life Technologies) according to the manufacturer’s protocol, and phagocytosis assays performed 72 hours later.

Peritoneous macrophages were collected in 5mL of ice-cold sterile 1X PBS supplemented with antibiotics that were introduced in the peritoneum of euthanized mice and retrieved with a 26G needle after gentle shaking (Enderlin et al., 2024). After centrifugation at 300g for 10min, macrophage pellets were resuspended in RPMI with antibiotics (100U/mL penicillin and 100μg/mL streptomycin) (all from Gibco), cells were counted and seeded at a density of 40,000 cells per well on assay cartridges and 120,000 cells per well for 96-well plates. After 2 hours of adhesion, unattached cells were washed out, new medium was added and macrophages were cultured in a humidified atmosphere containing 5% CO_2_ at 37°C for 24 to 72 hours.

### Phagocytosis Assay

To assess their phagocytic capacity cells were challenged with approximately 10 POS/cell resuspended in DMEM (Parinot et al., 2014) for 1.5 and 3 hours. In some assays, the ER stress inducer tunicamycin (Sigma) was added to the cells at a sub-lethal concentration (0.062μg/mL) 24 hours before the assay. At the end of the incubation, cells were washed 3 times with PBS-CM (1X PBS, 0.2mM Ca^2+^, 0.1mM Mg^2+^). For each condition half of the wells were treated with trypan blue (Life Technologies) for 10min to quench the fluorescence of surface-bound FITC-labeled POS and thus quantify only internalized POS (Finnemann et al., 1997). After 2 PBS-CM washes of trypan blue wells, all cells were fixed with ice-cold methanol and nuclei were counterstained with 1mg/mL DAPI in 1X PBS (FluoProbes). FITC-POS and DAPI-labeled nuclei signals were quantified using a fluorescence microplate reader (Infinite M1000, Magellan 6 software, Tecan).

### Statistical analysis

Shown immunoblot and immunofluorescence data are representative of at least 3 independent experiments. Values are given as the mean ± standard deviation (s.d.) of at least 3 independent experiments as indicated in each figure legend. Statistical significance was determined using *t* tests corrected for multiple comparisons using the Holm-Sidak method using the GraphPad Prism software. Differences were considered significant according to the following criteria: * *P* < 0.05, ** *P* < 0.01, *** *P* < 0.001 and **** *P* < 0.0001.

## Author contributions

E.F.N. designed the study; E.F.N, F.H., E.M.V., Q.R., J.E., N.H. and D.L. performed experiments and collected the data; E.F.N., F.H. and E.M.V. analyzed and interpreted the data; A.P. helped with the microplate reader program set-up and related data acquisition; G.M.P. and T.L. provided expertise with the metabolic assays and helped with the manuscript; E.F.N., A.H. and E.M.V. prepared the manuscript. All authors have read and agreed to the published version of the manuscript.

## Acknowledgements

The authors would like to thank Amélie Lacombe and Olivier Brégerie from the PreclinICAN core facility (Institute of Cardiometabolism And Nutrition, Sorbonne Université) for the metabolic phenotyping of our mice in calorimetric cages.

This work was supported by the Laboratory of Excellence Program (LABEX) [LIFESENSES: ANR-10-LABX-65, Project Grant to E.F.N.] and IHU FOReSIGHT [ANR-18-IAHU-0001, Project Grant to E.F.N.] from Agence Nationale de la Recherche, by the joint Funding Program from Union Nationale des Aveugles et Déficients Visuels (UNADEV)-Alliance pour les Sciences de la Vie et de la Santé (AVIESAN) [Project Grant to EFN], by Fondation de France [“Allocation Jeunes Chercheurs en Ophtalmologie” PhD funding to E.M.V.], and by Centre National de la Recherche Scientifique (CNRS). Additionally, the Institut de la Vision is funded by Institut National de la Santé et de la Recherche Médicale (INSERM), Sorbonne Université and CNRS, and is affiliated to DIM C-BRAINS, funded by the Conseil Régional d’Ile-de-France. The authors declare no competing financial interests.

## Abbreviations

1-DG: 2-deoxy-D-glucose;
4-HNE: 4-hydroxynonenal;
adRP: autosomic dominant RP;
ATF6: activating transcription factor 6;
ATP: adenosine triphosphate;
ATP6: ATP synthase 6;
ATPp: ATP production;
BR: basal respiration;
BHI: bioenergetic health index;
Bip: immunoglobulin heavy chain binding protein;
BPTES: glutaminase Inhibitor II,Bis-2-(5-phenylacetamido-1,2,4-thiadiazol-2-yl)ethyl sulfide;
Chop: C/EBP-homologous protein;
CoxIV: cytochrome C oxidase subunit 4;
ER: endoplasmic reticulum;
ECAR: extracellular acidification rate;
FCCP: carbonyl cyanide-p-trifluoromethoxyphenylhydrazone;
Grp78: glucose-regulated protein 78;
GlycoPER: glycolytic proton efflux rate;
HBSS: Hank’s balanced salt solution;
IRE1α: inositol requiring enzyme-1 alpha;
iPSC-RPE: RPE cells derived from induced pluripotent stem cells;
ND4: NADH-ubiquinone oxidoreductase chain 4;
OXPHOS: oxidative phosphorylation;
OCR: oxygen consumption rate;
PERK: protein kinase R-like endoplasmic reticulum kinase;
POS: photoreceptor outer segments;
PRPF: pre-mRNA processing factor;
ROS: reactive oxygen species;
RPE: retinal pigment epithelial;
RP: retinitis pigmentosa;
shRNA: short hairpin RNA;
siRNA: small interfering RNA;
SOD: superoxide dismutase;
tri-snRNP: tri-small nuclear ribonucleoprotein particle;
UPR: unfolded protein response;
wt: wildtype.

## Notes

### Competing Interest Statement

The authors have declared no competing interest.

