## Supplementary materials for "Energetic failure and oxidative stress underlie the *Prpf31* splicing factor-related mouse phenotype"

**Supplemental materials**

**Supplementary Table 1.** Name, reference, species and dilutions corresponding to the various antibodies used in immunohistochemical and western blot experiments.

| *Antibody* | *Reference* | *Species* | *IHC dilution* | *WB dilution* |
|---|---|---|---|---|
| 4-Hydroxynonenal | Novus Biologicals NB100-63093 | Goat | 1:200 | |
| 8-Oxoguanine (483.15) | Millipore MAB3560 | Mouse mAb | 1:200 | |
| ATF6 (70B1413.1, Nter) | Novus Biologicals NBP1-40256 | Mouse mAb | | 1:1,000 |
| ATP synthase (7H10BD4F9) | Life technologies 459240 | Mouse mAb | | 1:2,000 |
| Catalase (CAT-505) | Sigma C0979 | Mouse mAb | | 1:2,000 |
| CHOP (L63F7) | Cell Signaling 2895 | Mouse mAb | | 1:1,000 |
| COXIV | Biorbyt orb382539 | Rabbit | | 1:1,000 |
| eIF2α (D7D3) | Cell Signaling 5324 | Rabbit mAb | | 1:1,000 |
| P-eIF2α (119A11) | Cell signaling 3597 | Rabbit mAb | | 1:1,000 |
| ND4 | Biorbyt orb374304 | Rabbit | | 1:1,000 |
| Nitrotyrosine | Millipore 06-284 | Rabbit | 1:200 | |
| NPCP (Mab414) | Abcam ab24609 | Mouse mAb | | 1:2,000 |
| SOD1 | Biorbyt orb27559 | Rabbit | | 1:2,000 |
| SOD2 | Biorbyt orb76289 | Rabbit | | 1:1,000 |
| SOD3 | R&D Systems AF4817 | Goat | | 1:1,000 |
| Tubuline (TU-01) | Novus Biologicals NB500-333 | Mouse mAb | | 1:5,000 |

**Supplementary Table 2.** Mouse gene names and corresponding forward (F) and reverse (R) primer sequences.

| ***Gene*** | ***5'-3' Sequence*** | |
|---|---|---|
| Mouse *Atf4* | F | CGAGATGAGCTTCCTGAACAGC |
| | R | GGAAAAGGCATCCTCCTTGC |
| Mouse *Atf6* | F | TACCACCCACAACAAGACCA |
| | R | TGATGATCCCGGAGATAAGG |
| Mouse *ATP6* | F | CGTAATTACAGGCTTCCGACA |
| | R | AGCTGTAAGCCGGACTGCTA |
| Mouse *Bip/Grp78* | F | TGTGGTACCCACCAAGAAGTC |
| | R | TTCAGCTGTCACTCGGAGAAT |
| Mouse *Catalase* | F | CCACAGTCGCTGGAGAGTCA |
| | R | GTTTCCCACAAGGTCCCAGTT |
| Mouse *Chop* | F | CCAACAGAGGTCACACGCAC |
| | R | TGACTGGAATCTGGAGAGCGA |
| Mouse *CoxIV* | F | GGCTGCGTCTTCTTCTTCAT |
| | R | ATGGGGTTGCTCTTCATGTC |
| Mouse *Nd4* | F | CAATCTGCTTACGCCAAACA |
| | R | GTGATGATGTGAGGCCATGT |
| Mouse *RPLP0* | F | CCTGAAGTGCTCGACATCAC |
| | R | TGCCAGGACGCGCTTGTAC |
| Mouse *SOD1* | F | AGGGAACCATCCACTTCGAG |
| | R | AAAATGAGGTCCTGCACTGG |
| Mouse *SOD2* | F | GGCCAAGGGAGATGTTACAA |
| | R | GCTTGATAGCCTCCAGCAAC |
| Mouse *SOD3* | F | GACACCGTTCCTCTGTGTCC |
| | R | CAAAGACTAGGGGCTTGTGG |

### Supplementary Figure 1

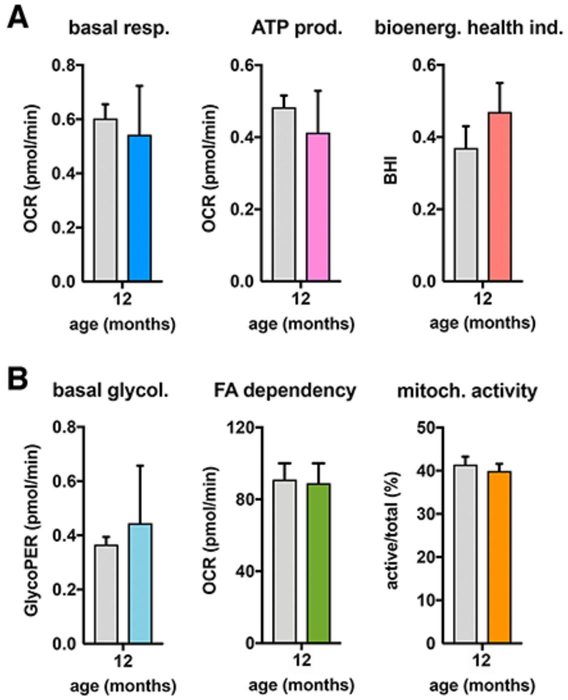


**Figure S1.** Normal mitochondrial function, glycolysis and fatty acids dependency in peritoneal macrophages from *Prpf31*$^{+/-}$ mice. **(A)** Basal respiration (left, blue), ATP production (middle, pink) and bioenergetic health index (right, dark orange) profiles of 12-month-old wildtype control (grey bars) and *Prpf31*$^{+/-}$ (colored bars) peritoneal macrophages. All are expressed as OCR measures in pmol/min. **(B)** Basal glycolysis (left, light blue), fatty acids dependency (FA, middle, green) and mitochondrial activity (right, light orange) profiles of 12-month-old wildtype control (grey bars) and *Prpf31*$^{+/-}$ (colored bars) peritoneal macrophages. Glycolysis is expressed as GlycoPER measures in pmol/min, FA dependency as OCR measures in pmol/min and mitochondrial activity as the percentage (%) of active mitochondria as indicated. Mean ± s.d., n = 3 (A) and n = 3-6 (B), multiple *t* test, all non significant.

**Supplementary Figure 2**

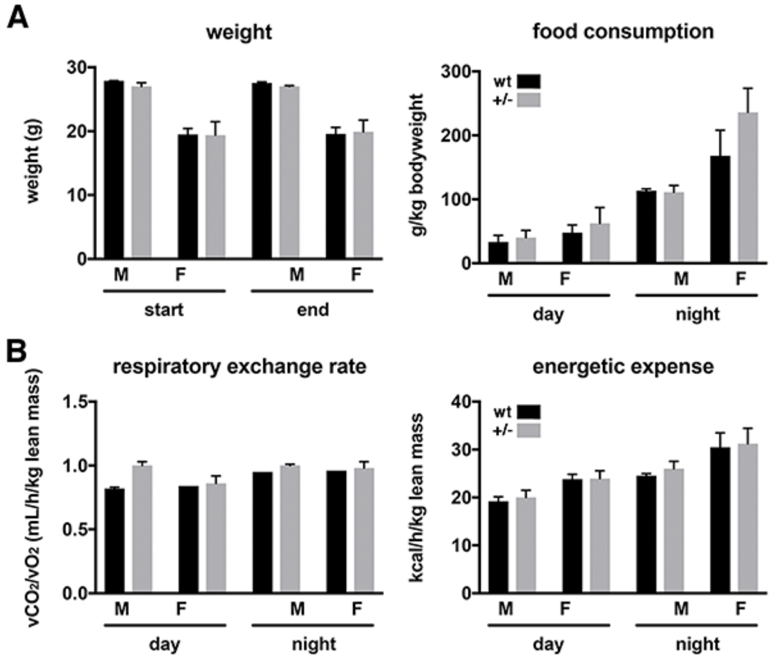


**Figure S2.** Normal body metabolism in *Prpf31*$^{+/-}$ mice. **(A)** Weight (left, g) and food consumption (right, g/kg body weight) measures in 3-month-old wildtype control (black bars) and *Prpf31*$^{+/-}$ (grey bars). Weight was measured at the beginning and the end of the 1-week follow-up period, and food consumption was monitored during the day and nigt shift periods. **(B)** Day and night respiratory exchange rate (left, $vCO_2/vO_2$ in mL/h/kg lean mass) and energetic expense (right, kcal/h/kg lean mass) measures in 3-month-old wildtype control (black bars) and *Prpf31*$^{+/-}$ (grey bars). Mean ± s.d., n = 4 females and 2 males of each genotype, all non significant.
